# Extending the Acute Skin Response Spectrum to Include the Far-UVC

**DOI:** 10.1101/2024.07.19.604306

**Authors:** Natalia E. Gutierrez-Bayona, Camryn Petersen, Raabia H. Hashmi, Manuela Buonanno, Igor Shuryak, Brian Ponnaiya, Norman J. Kleiman, David J. Brenner, David Welch

## Abstract

Guidance on maximal limits for ultraviolet (UV) exposure has been developed by national and international organizations to protect against adverse effects on human skin and eyes. These guidelines consider the risk of both acute effects (i.e. erythema and photokeratitis) and delayed effects (e.g., skin and ocular cancers) when determining exposure limits, and specify the dose a person can safely receive during an 8-hour period without harmful effects. The determination of these exposure limits relies on the action spectra of photobiological responses triggered by UV radiation that quantify the effectiveness of each wavelength at eliciting each of these effects. With growing interest in using far-UVC (200-235 nm) radiation to control the spread of airborne pathogens, recent arguments have emerged about revisiting exposure limits for UV wavelengths. However, the standard erythema action spectra, which provides some of the quantitative basis for these limits, has not been extended below 240 nm. This study assists to expand the erythema action spectrum to far-UVC wavelengths using a hairless albino mice model. We estimate that inducing acute effects on mouse skin with 222 nm radiation requires a dose of 1,162 mJ/cm^2^, well above the current ACGIH skin exposure limit of 480 mJ/cm^2^.

## INTRODUCTION

Exposure of human skin to ultraviolet (UV) radiation can cause an array of phototoxic effects including induction of inflammatory responses that can lead to skin ageing^1^, sunburns (erythema),^2^ and skin cancer.^3,4^ The UV light spectrum spans wavelengths from 100-400 nm and can be subdivided into three bands: UVA (315-400 nm), UVB (280-315 nm), and UVC (100-280 nm).^5^ Contained within the UVC band is a sub-band known as far-UVC that ranges from 200 - 235 nm. Each of these UV bands, characterized by their specific wavelengths and photon energy, are known to exert a variety of effects on the skin due to their penetrative capacity and interaction with different biophysical targets found within cells. The longest wavelengths, UVA, can penetrate deep into the dermis due to their lower photon energies that do not get absorbed by cellular biomolecules but can nevertheless induce the formation of damaging reactive oxygen species.^6^ In contrast, shorter wavelengths found within the UVB and UVC bands are mostly absorbed in the epidermis by interactions with major cellular biomolecules such as DNA and proteins. ^2,7,8^

The first line of defense of the skin against UV radiation is the outermost layer of the epidermis known as the stratum corneum (SC). Acting as a biomechanical barrier, the SC provides protection against various environmental factors and serves as an optical barrier against UV light.^7^ Research indicates that the SC absorbs UVC light most effectively, resulting in more significant mechanical and structural damage to the SC, than that induced by UVB and far less than that from UVA exposure.^7,9^ The transmission and thickness of the stratum corneum determines the level of optical protection it provides to the dividing cells found in the underlying skin layers.^10–12^ The high absorption of UVC light by the SC, particularly in the far-UVC region, is attributed to absorption by peptide bonds of proteins and lipids, while absorption in the UVB range is mainly due to the protein keratin and in the UVA range by melanin.^8,13–15^

Due to the different harmful effects that each component of the UV spectrum has on the skin and eyes,^5^ guidance on maximal safety limits for UV exposure has been developed by national and international groups.^16,17^ These guidelines were established to minimize both the risk of acute adverse effects (i.e. erythema and photokeratitis),^18^ as well as delayed effects (i.e. skin and ocular cancers).^19,20^ These exposure limits depend on the wavelength and the photobiological responses induced, where thresholds for each biological effect vary strongly with wavelength. To this extent, action spectra have been developed that allows quantitative measure of the spectral effectiveness of each wavelength at eliciting specific biological effects.^21^

The earliest and most widely used guidelines for human exposure limits to radiation are issued by the American Conference of Governmental Industrial Hygienists (ACGIH) and are referred to as Threshold Limit Values (TLVs).^22^ TLVs have been defined as the dose that a person can be safely exposed to in a 8 h period without developing skin or eye injuries.^22,23^ Similar guidelines on the limits of exposure have been implemented by the International Commission on Non-Ionizing Radiation Protection (ICNIRP) in collaboration with the World Health Organization (WHO).^17^

In 2021, a review by Sliney and Stuck argued for raising the exposure limits for wavelengths below 250 nm.^24^ They based their argument on numerous studies that found that the existing thresholds set for 222 nm and 207 nm to be overly conservative considering the minimal biological effects induced by these wavelengths.^24–31^ Subsequently, in 2022, ACGIH increased the TLVs for wavelengths below 240 nm and established separate limits for eyes and skin at wavelengths below 300 nm. Notably, higher exposure limits were set for wavelengths below 300 nm for cases where human skin is exposed but the eyes are protected.^32^ The TLVs for the shorter wavelengths in the far-UVC region were increased for both eye and skin exposures. Radiation at 222 nm, which is of particular interest for its antimicrobial properties within occupied spaces using filtered KrCl excimer lamps, ^33,34^ had the TLVs raised from 23 mJ/cm^2^ to 160 mJ/cm^2^ for eyes and to 480 mJ/cm^2^ for the skin. ^24,32,35^

Even though considerable evidence suggests the new skin threshold limits for wavelengths below 240 nm pose minimal risk,^19,24^ the standard erythema action spectra, which provide the quantitative basis for these limits, have not been extended below 240 nm.^16,36,37^ Therefore, there is a need to expand the erythema action spectrum curve to include results down to 200 nm to better define the potential hazards of far-UVC exposures.^38–40^ Previous research on murine models regarding far-UVC skin safety noted that erythema may not be the most accurate indicator of acute skin response at these wavelengths, as scale formation (flaking) has been observed to occur in the absence of erythema following 222 nm UV exposure.^27,38,41^ Consequently, more appropriate evaluation methods have been adopted to determine the acute effects of far-UVC on the skin. These methods include a new threshold dose estimate termed the minimal perceptible response dose (MPRD), which considers the development of erythema, edema, scales, and fissures, rather than solely relying on the minimal erythema dose (MED). It has been observed that using MPRD evaluations to assess the effects of UVB radiation provides action spectra similar to the human erythema action spectra ^38^. In the current study, we expand the erythema response spectrum down to 200 nm in hairless, albino, SKH-1 mice using MPRD evaluations. The SKH-1 experimental mouse model is often used in dermatological UVB exposure studies because this strain is particularly sensitive to UV effects and develops skin damage similar to that seen in humans.^40^ Irradiations were performed with wavelengths between 200 and 270 nm in 5 nm increments using a UV monochromator with a narrow bandwidth emission.^24,42^ The use of a narrow bandwidth provided more detailed information about the sharp increase in erythema response observed in the 230-240 nm range. This study reports minimal perceptible response threshold doses for skin for wavelengths across the UVC spectrum.

## MATERIALS AND METHODS

### Monochromatic source

An optical system was assembled to deliver monochromatic UV light to specific locations on the dorsal surface of mice. An EQ-77 high-brightness, long-life, broadband laser-driven light source (Energetiq Technology, Inc., Wilmington, MA) was focused into a Newport Cornerstone 260 1/4 m monochromator (CS260-RG-2-FH-A, Newport, Irvine, CA). The monochromator was equipped with a 1201.6 g/mm plane blazed holographic reflection grating (#53-*-200H with master no. 5482; 250 nm nominal blaze wavelength, Newport) and a 600 g/mm plane holographic grating (#53-*-313H with master no. 6018, 250 nm nominal blaze wavelength, Newport) to maximize optical throughput in the UVC. Fixed slits with a slit size of 600 μm (77216, Newport) were used for all experiments. With this setup, the monochromator output bandwidth at full width half maximum (FWHM) was nominally 1.9 nm for the 1201.6 g/mm grating and 3.8 nm for the 600 g/mm grating.

The optical power was measured with an 818-UV/DB low-power UV enhanced silicon photodetector (Newport). The average size of the exposure area was measured to be 0.23 cm^2^, which was determined by exposing radiation sensitive film (OrthoChromic Film OC-1, Orthochrome Inc., Hillsborough, NJ), scanning the film using an Epson Perfection V850 Pro flatbed scanner (Epson America, Los Alamitos, CA), and determining the exposed area using ImageJ.^43^ The irradiance was determined by dividing the output power for a chosen wavelength by the exposure area. The spectral irradiance was measured with a Gigahertz-Optik BTS2048-UV Spectroradiometer (Gigahertz-Optik, Inc., Amesbury,MA). The FWHM for the nominally 1.9 nm setup was measured as 2.2 nm, and the FWHM for the nominally 3.8 nm setup was 4.4 nm. These two exposure scenarios are referred to as the 2 nm or 4 nm bandwidth exposures throughout this manuscript.

### Mice

Eight-week-old male albino hairless SKH1-Elite mice, strain 477, were obtained from Charles River Laboratories (Wilmington, MA). Mice were allowed to acclimatize for one to two weeks after receipt. They were housed in standard filter top cages with *ad libitum* access to water and Purina laboratory chow. Just prior to irradiation, the 8-9 week old mice were anesthetized with an intraperitoneal injection of a ketamine/xylazine mixture (100 mg/kg /10 mg/kg respectively). If an animal awakened during irradiation, an additional injection of 0.125 mL of the ketamine/xylazine mixture was administered. The depth of anesthesia was confirmed by toe pinch. Puralube ophthalmic ointment (Dechra Veterinary Products, Overland Park, KS) was applied to the eyes during anesthesia. A grid of approximately 2 cm x 4 cm was drawn on the mouse dorsal skin with a permanent marker to define exposure regions and the dorsal side of the mouse was photographed prior to any exposure (Figure 1a). Pictures of the mouse dorsal skin were taken with a Canon EOS Rebel T7 (Canon USA, Huntington, NY). The camera was mounted with its lens positioned 22 cm from the surface where the mouse was located, positioned between two light boxes. All animal procedures were approved by the Columbia University Institutional Animal Care and Use Committee (IACUC).

**Figure 1.**
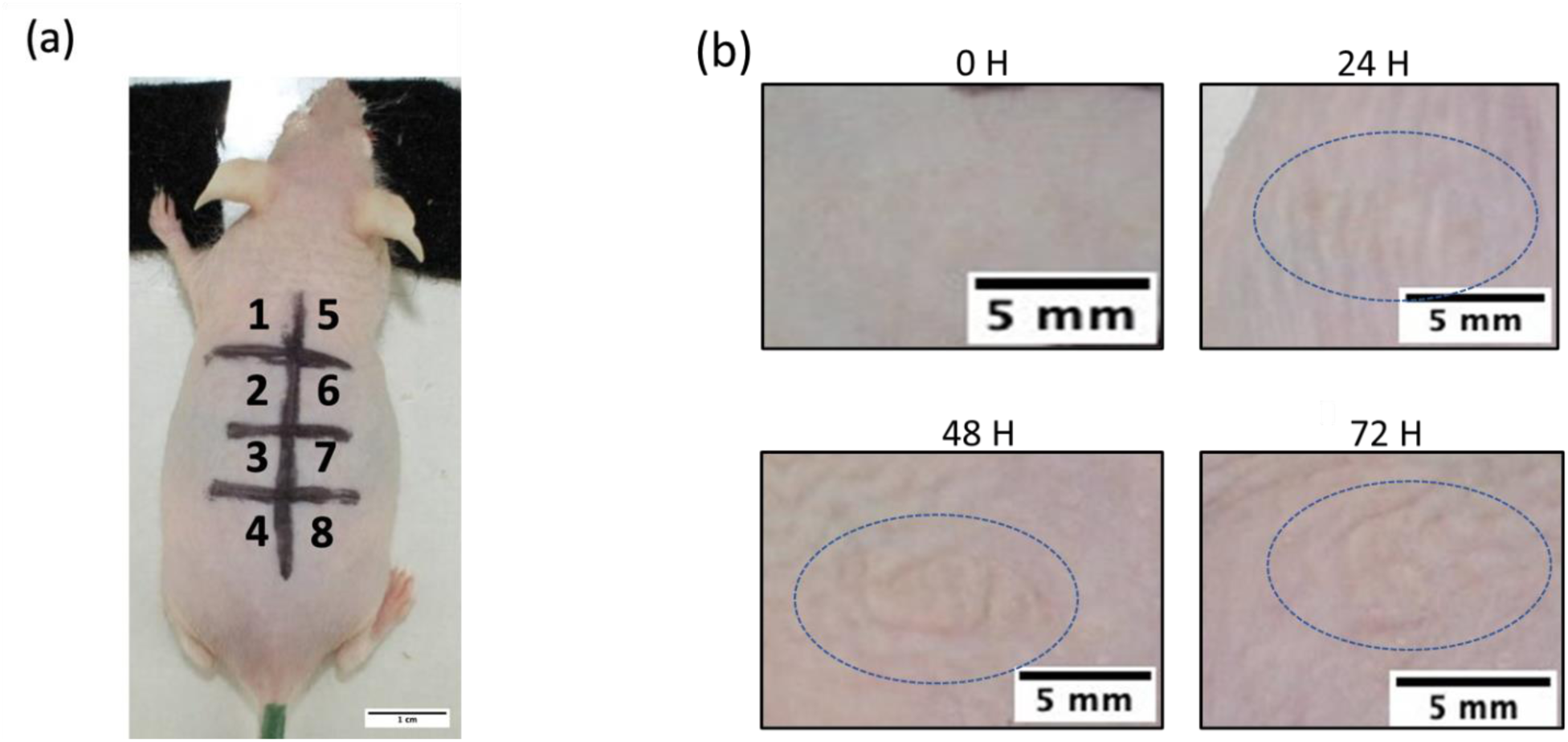
Hairless mice skin irradiated with monochromatic UV light based on an eight-section grid drawn on the dorsal side of the animal (a) 2 cm x 4 cm grid divided into eight 1 x 1 cm sections that were randomly selected for irradiations. (b) Images representing the progression of general skin damage observed on the dorsal skin of mice over a 72-hour period. The adverse effects delimited by the oval shape include scaling and change in color of the skin. The pictures are from skin spots exposed to 1120 mJ/cm^2^ from 225 nm.

### Experimental Design

To estimate the dose that elicited the minimal perceptible response for each wavelength from 200 nm to 270 nm in 5 nm steps, we utilized a two-stage process. The first stage exposed a single mouse to six different doses at a single wavelength to roughly determine the dose that induced the minimal perceptible skin response. To determine the exposure doses for each mouse in this first stage, the current 8-hour exposure limit as defined by ICNIRP for each respective wavelength, was used as well as 2, 4, 8, 16, and 32 times the ICNIRP exposure limit (Table S1).^17^ The regions that were exposed within the grid drawn on the mouse dorsal skin (Figure 1a) were randomly selected for each mouse and the remaining two regions on the grid were left as unexposed controls. The first step exposures were performed with a bandwidth of 2 nm, except for the 200 nm exposures, which used a 4 nm FWHM (Figure 2) due to the large doses required and the limited time that the mouse could be maintained under anesthesia. The high doses for 200 nm, 205 nm, and 210 nm required exposure times that exceeded the allowable mouse anesthesia time; therefore the 6 exposures were split across 2 mice instead of a single mouse.

**Figure 2.**
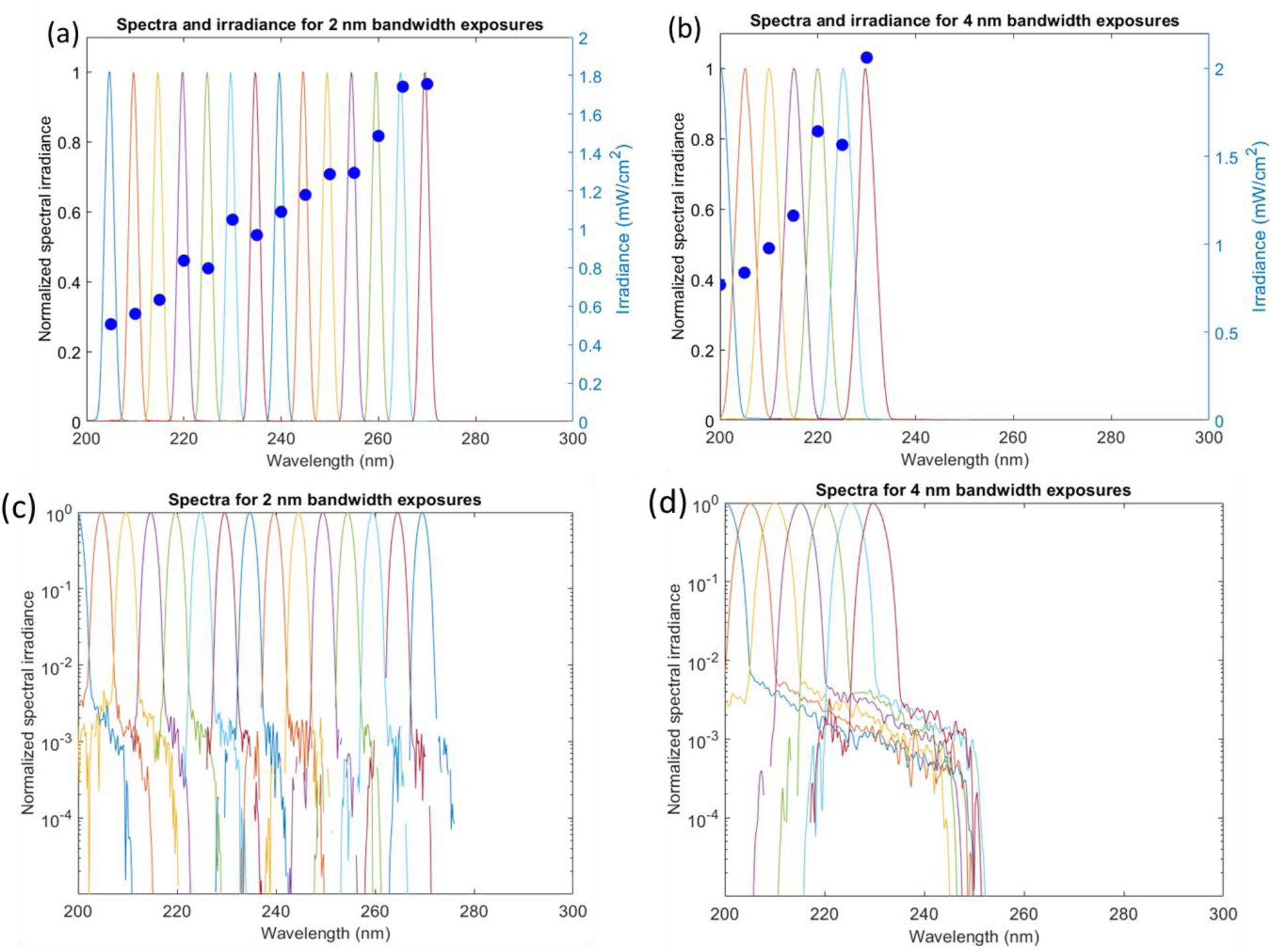
Measured spectra and irradiance values for 2 nm (235 – 270 nm) and 4 nm (200 – 230 nm) bandwidth exposures. (a-b) Spectra and irradiance value used for exposures at each wavelength plotted on a linear scale (c-d) Spectra plotted on a logarithmic scale showing the amount of stray radiation found at each wavelength.

Mice were evaluated at 24 h, 48 h, and 72 h after irradiation. Each skin spot was evaluated for the presence or absence of erythema, edema, flaking, or fissuring by two team members who were blinded to the dose. Additionally, a photograph of the dorsal skin of each mouse was taken at each evaluation time. The highest dose that was administered for each wavelength that was determined to be negative for all the test criteria was assigned to be the new estimated safe dose for that wavelength.

Following the establishment of a roughly estimated safe dose for each wavelength during the first stage of the process (20 mice used), the second stage made use of cohorts of 8 mice that helped to further narrow the threshold estimate (123 mice in total). For each wavelength, all 8 mice were exposed to a set of doses including the newly established estimated safe dose at each respective wavelength followed by 1.25x, 1.5x, 1.75x, and 2x this new estimated safe dose (Table S1). The 5 exposure sites were randomly assigned to the 8 positions at the dorsal of the mouse, with the remaining 3 positions used as unexposed controls. Exposures for wavelengths from 235 nm to 270 nm were performed with a bandwidth of 2 nm (Figure 2a). For wavelengths of 230 nm and lower, the exposure bandwidth was increased to 4 nm (Figure 2b), as the new dose estimates established during the first stage indicated very high doses were required. This increase in the exposure bandwidth increased the optical power throughput of the monochromator system, shortening the exposure time for a given dose. Even with the increased bandwidth, the large doses determined for wavelengths of 215 nm and below required a reduction in the number of exposures to each mouse to allow for the total exposure time to remain within the time the mouse could be kept anesthetized. For 215 nm the exposures were limited to 1x, 1.25, and 1.5x the estimated dose; for 210 nm and 205 nm the exposures were limited to 1x and 1.25x the estimated dose; and for 200 nm the exposure was limited to the single dose.

### Data Analysis

From the collected data, logistic regression models were constructed and compared using the *glmulti R* package. Various combinations of predictor terms, including wavelength (*w*, in nm), *w^2^*, log_10_-transformed dose (*D*, mJ/cm^2^; it was set to 0 if the original dose was 0), *D^2^*, evaluation time point (*t*, hours), and *t^2^*, were explored. Additionally, multiplicative interactions between these terms were examined. The selection of the most appropriate model variant was based on the Akaike information criterion with sample size correction (AIC). To account for potential variability between individual mice and for correlated responses within the same mouse, random effects were incorporated by considering mouse and cage effects for both intercepts (baselines) and “slope” terms in the best-supported model. The best supported mixed effects model was also selected based on AIC.

The dose value at which the predicted probability of the effect (MPRD) would reach 50% (D_50_) was calculated for the best-supported mixed effects model using its best-fit parameters. Uncertainties of D_50_ for each wavelength were calculated using refitting of the best-supported model to 1,000 bootstrapped versions of the data set. Confidence intervals (95% CIs) for D_50_ were estimated from the bootstrapped data fits using the 2.5^th^ and 97.5^th^ percentiles of the D_50_ distribution at each wavelength.

## RESULTS

### Minimal Skin Reactions to UVC Exposures

Similar to previous studies on acute skin responses to UV radiation,^38,41^ mouse skin was visually evaluated for the obvious development of edema, scaling, erythema, and/or fissures 24-, 48-, and 72-hours after irradiation (Figure 1b). No grading was made to quantify the severity of the skin reactions. Across all exposure doses and wavelengths tested, a wide range of sensitivity was observed within the mouse population (Figure 3), with no discernible trend in skin reactions. However, erythema was the most consistently observed and easily perceived endpoint for the highest exposure doses used at wavelengths above 215 nm (Figure 1b). For some of the mice population, scale formation (scaling) on the irradiated portion of the skin was also observed, primarily developing at the 48-h mark or later for wavelengths above 215 nm.

**Figure 3.**
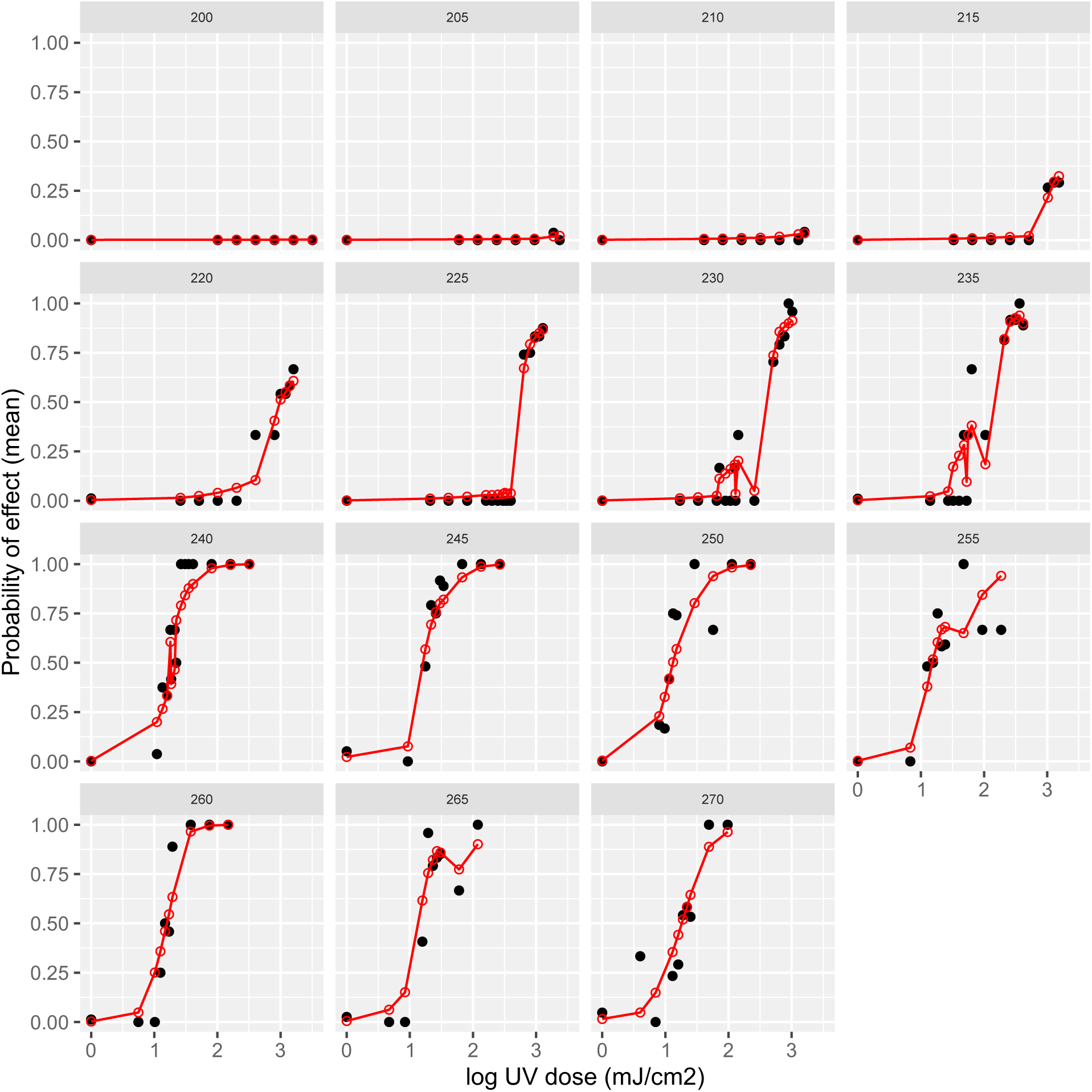
Comparison of the probability effects predicted by the best-supported model (red) to the mean of the acute skin evaluations done throughout the study (black). The probability of a skin response is plotted for each dose at each wavelength tested. Each data point represents the exposures of multiple mice at one dose and the evaluations done at all times (24-, 48-, and 72-h) are combined.

The photographs taken of the skin reactions at each irradiation wavelength were found to be difficult to use for analysis of the MPRD. Upon reviewing the photographic records, it became apparent that some of the skin reactions could not be clearly differentiated due to the small size of the irradiated area. Based on visual inspections, erythema was the only skin reaction that was easily identifiable in the photographs (Figure 1b), while edema was visually discernible to a lesser extent. Scaling posed the most significant challenge to image, particularly at doses below the maximum exposure levels.

### Data Analysis

Time (24 h, 48 h, 72 h) of the evaluation for MPRD showed low importance and was ultimately discarded from the best-supported model structure. The best-supported mixed effects model had the following fixed effect’s structure, where *P* is the predicted effect probability, *D* is log_10_-transformed dose, *w* is wavelength, and *b_1_*-*b_3_* are adjustable parameters:

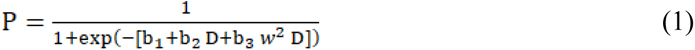

The probability was calculated for each exposure dose evaluated at every wavelength during the different stages of the study using the best-supported model, and then compared to the mean of the skin evaluation results (Figure 3). The slight differences observed in the fit of the predicted probability for each mouse were attributed to random effects. Once we confirmed the model accurately predicts the observed skin response according to dose and wavelength, we determined the dose at which the predicted probability of the skin effect would reach 50% (D_50_). We therefore solved the model equation (1) for *D*, where P=0.5:

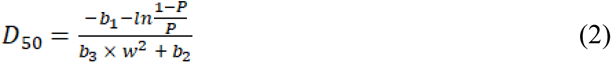

Notably, this D_50_ calculation is on a log_10_ scale, and generated using only the fixed effects terms from the model, ignoring inter-mouse variations. The observed skin effect values, reported on a linear scale, for wavelengths ranging from 200-215 nm (Table 1), never reached 50% (Figure 3), so the D_50_ values that were calculated for these wavelengths are very high (>10^4^) and extrapolations. The actual D_50_ could not be estimated in that wavelength region using the current data.

**Table 1.**
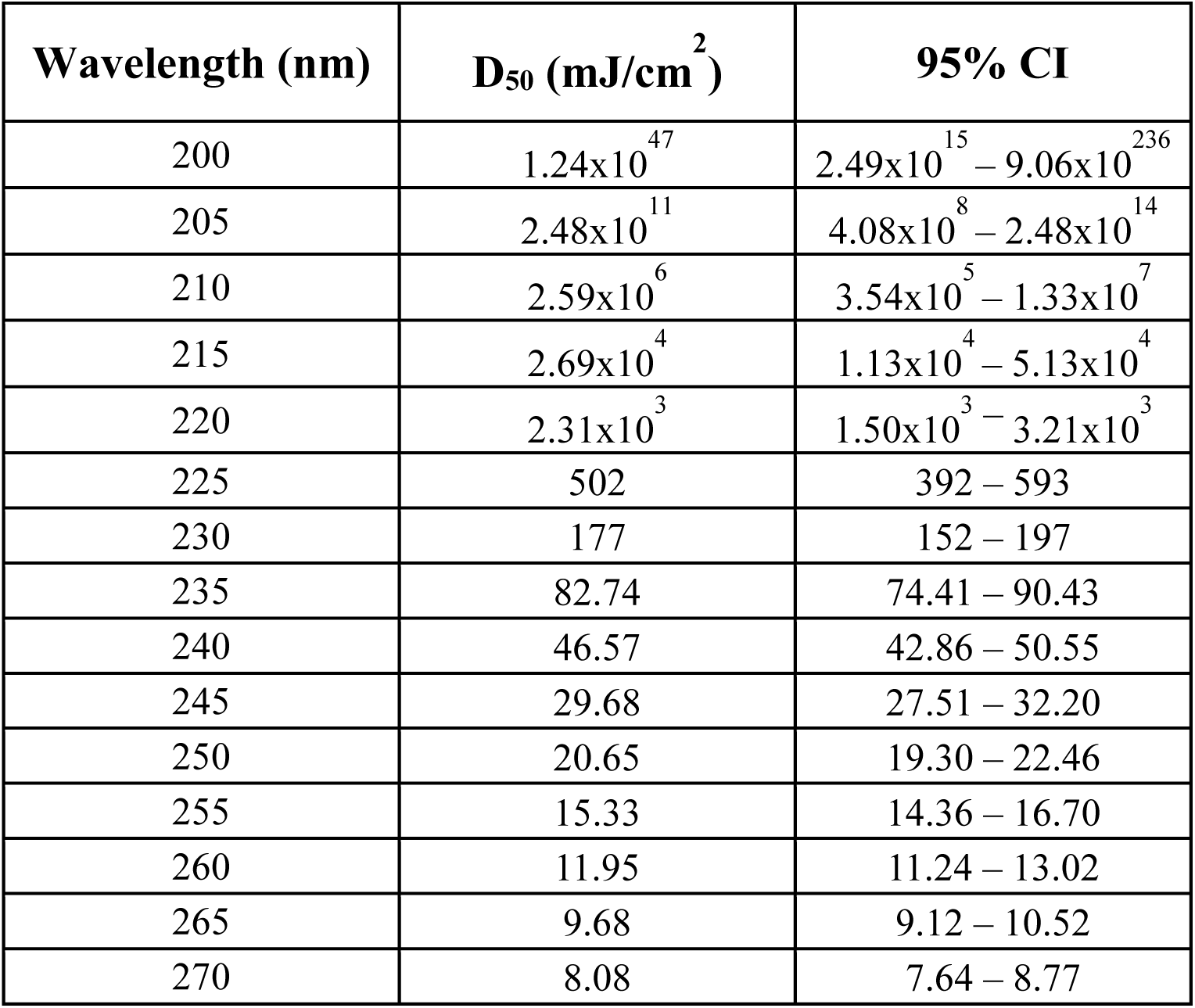
Doses (D_50_) at each wavelength that are predicted to induce a minimal perceptible skin response with a probability of 50%. Confidence intervals (95% CIs) were estimated from the bootstrapped data fits using the 2.5^th^ and 97.5^th^ percentiles of the D_50_ distribution at each wavelength.

| Wavelength (nm) | D<sub>50</sub> (mJ/cm<sup>2</sup>) | 95% CI |
| --- | --- | --- |
| 200 | 1.24x10<sup>47</sup> | 2.49x10<sup>15</sup> – 9.06x10<sup>236</sup> |
| 205 | 2.48x10<sup>11</sup> | 4.08x10<sup>8</sup> – 2.48x10<sup>14</sup> |
| 210 | 2.59x10<sup>6</sup> | 3.54x10<sup>5</sup> – 1.33x10<sup>7</sup> |
| 215 | 2.69x10<sup>4</sup> | 1.13x10<sup>4</sup> – 5.13x10<sup>4</sup> |
| 220 | 2.31x10<sup>3</sup> | 1.50x10<sup>3</sup> – 3.21x10<sup>3</sup> |
| 225 | 502 | 392 – 593 |
| 230 | 177 | 152 – 197 |
| 235 | 82.74 | 74.41 – 90.43 |
| 240 | 46.57 | 42.86 – 50.55 |
| 245 | 29.68 | 27.51 – 32.20 |
| 250 | 20.65 | 19.30 – 22.46 |
| 255 | 15.33 | 14.36 – 16.70 |
| 260 | 11.95 | 11.24 – 13.02 |
| 265 | 9.68 | 9.12 – 10.52 |
| 270 | 8.08 | 7.64 – 8.77 |

### Response Spectrum

The predicted threshold doses for acute skin reactions triggered by UVC wavelengths ranging from 200-270 nm revealed a sharp increase in the response spectra as far-UVC wavelengths decreased (Figure 4). This sharp increase becomes notable at 240 nm, further intensifying at lower wavelengths, indicating that higher doses are required in the lower wavelengths of the UVC spectrum to elicit an acute skin response compared to longer wavelengths. This suggests that the shorter wavelengths found within the far-UVC region of the spectrum are less efficient than longer wavelengths of UVC at triggering acute skin responses in mouse skin. Specifically, at the 222 nm wavelength, we predict that a dose of 1,162 mJ/cm^2^ is needed to induce an acute skin reaction response within 72 h, which is well above the current exposure skin limits set at 480 mJ/ cm^2^.^32^

**Figure 4.**
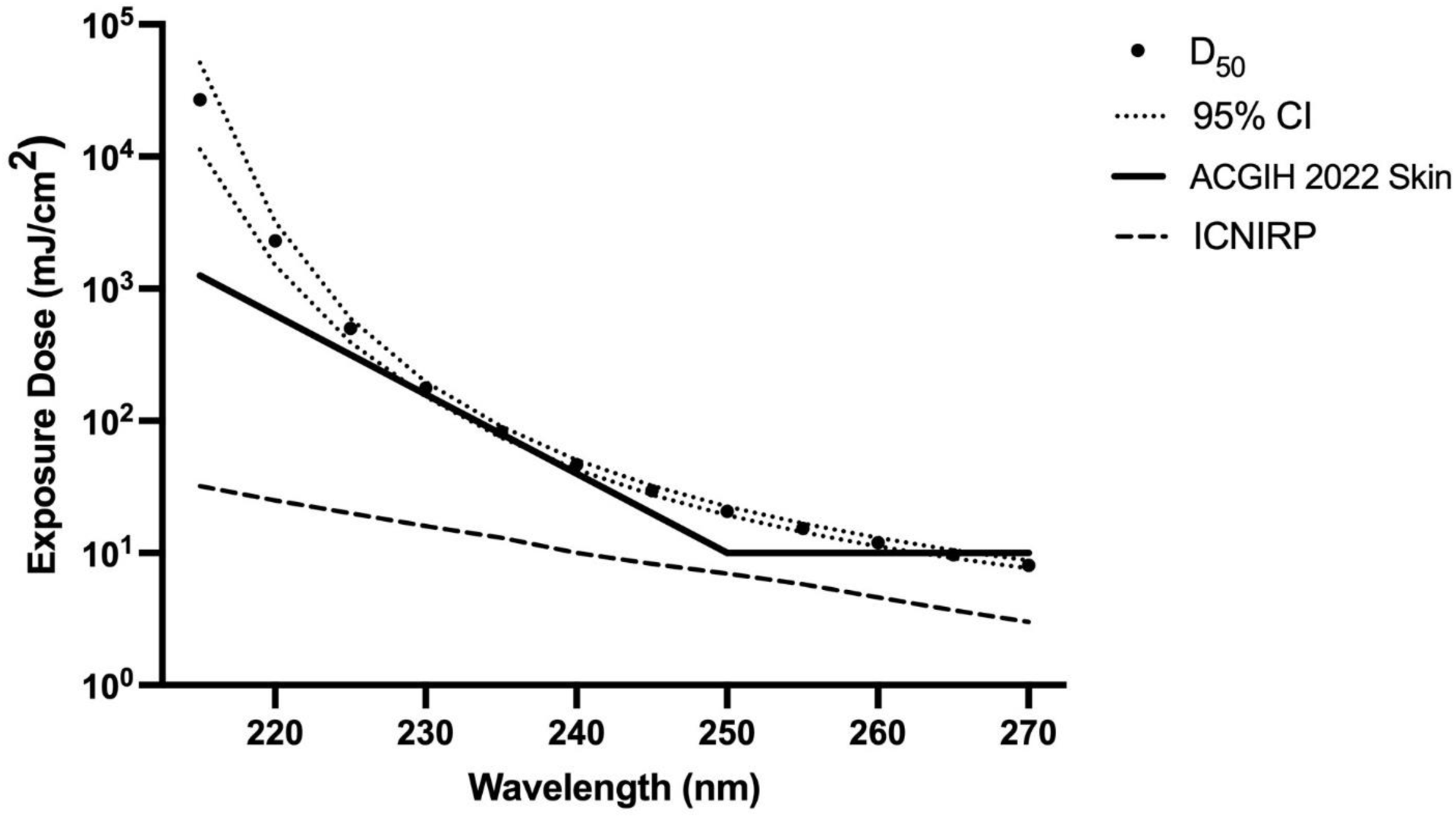
Acute skin response action spectra (predicted exposure dose vs wavelength) plotted in a linear scale. Estimated MPRD (D_50_) and the 95% confidence levels for wavelengths ranging from 215 to 270 nm compared to the UV hazard limits for the 2022 ACGIH skin TLVs and ICNIRP UV exposure limits. ^17,33^

## DISCUSSION

This study provides the most comprehensive analysis of acute skin response spectrum after UVC exposure (Figure 4). Previous reports of short-term skin responses to UV irradiation are limited for wavelengths below 250 nm primarily due to experimental inaccessibility to 1) a monochromator able to irradiate at spectral bandwidth less than 10 nm, and 2) limiting examinations to wavelengths from those principally emitted by gas lamps. ^24,42^ By taking proper consideration of the spectral bandwidth and utilizing narrow bandwidth exposures in 5 nm increments with minimal contributions of stray radiation (Figure 2), we are able to accurately report the effectiveness of UVC wavelengths at inducing short term skin effects down to 200 nm. This approach helps reduce the uncertainty inherent in predicting the effect of each wavelength on specific photobiological outcomes.^42^

The acute skin response spectrum reported in this study, which was derived from the predicted threshold doses (D_50_), revealed a sharp increase in response for the UVC wavelengths below 240 nm (Figure 4 & Table 1). The increase in the threshold doses needed to induce short-term skin effects have been attributed to the very short epidermal transmittance demonstrated by these short wavelengths in mammalian skin.^11,12^ It has been shown that wavelengths within the far-UVC spectrum become highly absorbed by stratum corneum significantly reducing their penetration into the deeper layers of the epidermis.^12^ The human stratum corneum has been determined to be typically 15-30 µm thick, while the epidermis has been found to range between 56-81 µm, varying depending on the body site.^44,45^ During this study, we measured the average thickness of the stratum corneum in hairless mice dorsal skin to be 4.70 ± 0.88 µm, while the epidermis measured 20.86 ± 5.52 µm (Table S2). This significant difference in stratum corneum and epidermis thickness between the SKH1 mouse model and human skin suggests that the skin responses observed in this study in hairless SKH1 mice, particularly for the far-UVC region,^7,12^ may underestimate the human skin UV threshold.^40,46^

In an earlier far-UVC safety skin study that used hairless SKH1 mice models,^41^ it was stated that a more appropriate consideration of acute effects for UVC radiation involves the development of erythema, edema, scales, and fissures, rather than solely relying on erythema. This approach was established by Yamano *et al*. ^41^ as a more accurate method to study the effects that far-UVC has on skin cutaneous responses, as erythema was rarely observed, while edema and scale formation were more evident. While their study used excimer lamps emitting wavelengths at 207, 222, 235 nm to explore the effects of far-UVC light on 1-cm square irradiated skin regions, our study irradiated a smaller area of mice skin. We found that erythema was the most easily perceptible skin response in static images among the proposed reactions (Figure 1b). The recognition of the formation of scales and edema proved to be challenging, as we only performed visual inspections, whereas Yamano *et al.*^41^ utilized both visual and palpation inspections to make these observations. Additionally, we noticed significant variability in the detectable skin reactions between mice (Figure 3), making it difficult to determine trends that could differentiate the effects triggered by the far-UVC region of the spectrum from other, longer wavelength, UVC radiation.

Based on the mouse skin acute response spectrum reported in this study, which covers UVC wavelengths from 200-270 nm (Figure 4), our data suggests that the revised skin TLVs established by ACGIH in 2022 for wavelengths below 230 nm is still overly conservative with respect to the risk of acute adverse skin effects. Specifically, we estimate that for wavelengths between 220-230 nm, the dose needed to induce a short-term effect on mouse skin ranges from 2,310 to 177 mJ/cm^2^ (Table 1), well above the current exposure skin limits set at 631 to 158.5 mJ/ cm^2^.^32^ While this mice skin acute threshold doses may not directly correspond to far-UVC safety in human skin, the findings of this study provides valuable data to guide development and planning of potential human acute skin exposure trials. In combination with studies to measure the effects of chronic far-UVC exposure, such as formation of UV-induced cyclobutane-pyrimidine dimers and 6-4 photoproducts, our data, and that of future studies, will inform the potential human health risks and acceptable threshold values for far-UVC radiation to enable its more widespread use as an antimicrobial technology.^19^

## ACKNOWLEDGMENTS

This work has been supported in part by CDC NIOSH Award #R01OH012312.

## SUPPLEMENTARY MATERIALS

Additional supporting information can be found online in the Supporting Information section at the end of this article.

Table S1 and Figure S2 can be found at DOI: 10.1562/2006-xxxxxx.s1.

## Notes

### Competing Interest Statement

The authors have declared no competing interest.

## REFERENCES

(1) Panich, U.; Sittithumcharee, G.; Rathviboon, N.; Jirawatnotai, S. Ultraviolet Radiation-Induced Skin Aging: The Role of DNA Damage and Oxidative Stress in Epidermal Stem Cell Damage Mediated Skin Aging. Stem Cells Int. 2016, 2016, e7370642. 10.1155/2016/7370642.

(2) D’Orazio, J.; Jarrett, S.; Amaro-Ortiz, A.; Scott, T. UV Radiation and the Skin. Int. J. Mol. Sci. 2013, 14 (6), 12222–12248. 10.3390/ijms140612222.

(3) Elwood, J. M.; Jopson, J. Melanoma and Sun Exposure: An Overview of Published Studies. Int. J. Cancer 1997, 73 (2), 198–203. 10.1002/(SICI)1097-0215(19971009)73:2<198::AID-IJC6>3.0.CO;2-R.

(4) Böni, R.; Schuster, C.; Nehrhoff, B.; Burg, G. Epidemiology of Skin Cancer.

(5) Diffey, B. L. Sources and Measurement of Ultraviolet Radiation. Methods 2002, 28 (1), 4–13. 10.1016/S1046-2023(02)00204-9.

(6) Cadet, J.; Douki, T.; Ravanat, J.-L. Oxidatively Generated Damage to Cellular DNA by UVB and UVA Radiation,. Photochem. Photobiol. 2015, 91 (1), 140–155. 10.1111/php.12368.

(7) Lipsky, Z. W.; German, G. K. Ultraviolet Light Degrades the Mechanical and Structural Properties of Human Stratum Corneum. J. Mech. Behav. Biomed. Mater. 2019, 100, 103391. 10.1016/j.jmbbm.2019.103391.

(8) Meinhardt, M.; Krebs, R.; Anders, A.; Heinrich, U.; Tronnier, H. Wavelength-Dependent Penetration Depths of Ultraviolet Radiation in Human Skin. J. Biomed. Opt. 2008, 13 (4), 044030. 10.1117/1.2957970.

(9) Diffey, B. L. A Mathematical Model for Ultraviolet Optics in Skin. Phys. Med. Biol. 1983, 28 (6), 647–657. 10.1088/0031-9155/28/6/005.

(10) Bruls, W. A. G.; Slaper, H.; Van Der Leun, J. C.; Berrens, L. Transmission of Human Epidermis and Stratum Corneum as a Function of Thickness in the Ultraviolet and Visible Wavelengths. Photochem. Photobiol. 1984, 40 (4), 485–494. 10.1111/j.1751-1097.1984.tb04622.x.

(11) Finlayson, L.; Barnard, I. R. M.; McMillan, L.; Ibbotson, S. H.; Brown, C. T. A.; Eadie, E.; Wood, K. Depth Penetration of Light into Skin as a Function of Wavelength from 200 to 1000 Nm. Photochem. Photobiol. 2022, 98 (4), 974–981. 10.1111/php.13550.

(12) Zamudio Díaz, D. F.; Klein, A. L.; Guttmann, M.; Zwicker, P.; Busch, L.; Kröger, M.; Klose, H.; Rohn, S.; Schleusener, J.; Meinke, M. C. Skin Optical Properties from 200 to 300 Nm Support Far UV-C Skin-Safety in Vivo. J. Photochem. Photobiol. B 2023, 247, 112784. 10.1016/j.jphotobiol.2023.112784.

(13) Bendit, E. G.; Ross, D. A Technique for Obtaining the Ultraviolet Absorption Spectrum of Solid Keratin. Appl. Spectrosc. 1961, 15 (4), 103–105. 10.1366/000370261774426957.

(14) Ou-Yang, H.; Stamatas, G.; Kollias, N. Spectral Responses of Melanin to Ultraviolet A Irradiation. J. Invest. Dermatol. 2004, 122 (2), 492–496. 10.1046/j.0022-202X.2004.22247.x.

(15) Anderson, R. R.; Parrish, J. A. The Optics of Human Skin. J. Invest. Dermatol. 1981, 77 (1), 13–19. 10.1111/1523-1747.ep12479191.

(16) Webb, A. R.; Slaper, H.; Koepke, P.; Schmalwieser, A. W. Know Your Standard: Clarifying the CIE Erythema Action Spectrum. Photochem. Photobiol. 2011, 87 (2), 483–486. 10.1111/j.1751-1097.2010.00871.x.

(17) The International Commission on Non-Ionizing Radiation Protection. GUIDELINES ON LIMITS OF EXPOSURE TO ULTRAVIOLET RADIATION OF WAVELENGTHS BETWEEN 180 Nm AND 400 Nm (INCOHERENT OPTICAL RADIATION). Health Phys. 2004, 87 (2), 171–186.

(18) Young, A. R. Acute Effects of UVR on Human Eyes and Skin. Prog. Biophys. Mol. Biol. 2006, 92 (1), 80–85. 10.1016/j.pbiomolbio.2006.02.005.

(19) Görlitz, M.; Justen, L.; Rochette, P. J.; Buonanno, M.; Welch, D.; Kleiman, N. J.; Eadie, E.; Kaidzu, S.; Bradshaw, W. J.; Javorsky, E.; Cridland, N.; Galor, A.; Guttmann, M.; Meinke, M. C.; Schleusener, J.; Jensen, P.; Söderberg, P.; Yamano, N.; Nishigori, C.; O’Mahoney, P.; Manstein, D.; Croft, R.; Cole, C.; de Gruijl, F. R.; Forbes, P. D.; Trokel, S.; Marshall, J.; Brenner, D. J.; Sliney, D.; Esvelt, K. Assessing the Safety of New Germicidal Far-UVC Technologies. Photochem. Photobiol. 00 (0), 1–20. 10.1111/php.13866.

(20) Yam, J. C. S.; Kwok, A. K. H. Ultraviolet Light and Ocular Diseases. Int. Ophthalmol. 2014, 34 (2), 383–400. 10.1007/s10792-013-9791-x.

(21) Erythema Reference Action Spectrum and Standard Erythema Dose; CIE S007E; Commission Internationale de l’Eclairage: CIE Central Bureau, Vienna, Austria, 1998.

(22) American Conference of Governmental Industrial Hygienists. Threshold Limit Values (TLVs) and Biological Exposure Index (BEIs) for Chemical Substances and Physical Agents for 2012; ACGIH, Cincinnati, OH, 2012.

(23) Sliney, D. Balancing the Risk of Eye Irritation from UV-C with Infection from Bioaerosols. Photochem. Photobiol. 2013, 89 (4), 770–776. 10.1111/php.12093.

(24) Sliney, D. H.; Stuck, B. E. A Need to Revise Human Exposure Limits for Ultraviolet UV-C Radiation†. Photochem. Photobiol. 2021, 97 (3), 485–492. 10.1111/php.13402.

(25) Buonanno, M.; Stanislauskas, M.; Ponnaiya, B.; Bigelow, A. W.; Randers-Pehrson, G.; Xu, Y.; Shuryak, I.; Smilenov, L.; Owens, D. M.; Brenner, D. J. 207-Nm UV Light—A Promising Tool for Safe Low-Cost Reduction of Surgical Site Infections. II: In-Vivo Safety Studies. PLoS ONE 2016, 11 (6), e0138418. 10.1371/journal.pone.0138418.

(26) Buonanno, M.; Ponnaiya, B.; Welch, D.; Stanislauskas, M.; Randers-Pehrson, G.; Smilenov, L.; Lowy, F. D.; Owens, D. M.; Brenner, D. J. Germicidal Efficacy and Mammalian Skin Safety of 222-Nm UV Light. Radiat. Res. 2017, 187 (4), 483–491. 10.1667/RR0010CC.1.

(27) Narita, K.; Asano, K.; Morimoto, Y.; Igarashi, T.; Nakane, A. Chronic Irradiation with 222-Nm UVC Light Induces Neither DNA Damage nor Epidermal Lesions in Mouse Skin, Even at High Doses. PLOS ONE 2018, 13 (7), e0201259. 10.1371/journal.pone.0201259.

(28) Yamano, N.; Kunisada, M.; Kaidzu, S.; Sugihara, K.; Nishiaki-Sawada, A.; Ohashi, H.; Yoshioka, A.; Igarashi, T.; Ohira, A.; Tanito, M.; Nishigori, C. Long-term Effects of 222-nm Ultraviolet Radiation C Sterilizing Lamps on Mice Susceptible to Ultraviolet Radiation. Photochem. Photobiol. 2020, 96 (4), 853–862. 10.1111/php.13269.

(29) Fukui, T.; Niikura, T.; Oda, T.; Kumabe, Y.; Ohashi, H.; Sasaki, M.; Igarashi, T.; Kunisada, M.; Yamano, N.; Oe, K.; Matsumoto, T.; Matsushita, T.; Hayashi, S.; Nishigori, C.; Kuroda, R. Exploratory Clinical Trial on the Safety and Bactericidal Effect of 222-Nm Ultraviolet C Irradiation in Healthy Humans. PLOS ONE 2020, 15 (8), e0235948. 10.1371/journal.pone.0235948.

(30) Kaidzu, S.; Sugihara, K.; Sasaki, M.; Nishiaki, A.; Igarashi, T.; Tanito, M. Evaluation of Acute Corneal Damage Induced by 222-Nm and 254-Nm Ultraviolet Light in Sprague– Dawley Rats. Free Radic. Res. 2019, 53 (6), 611–617. 10.1080/10715762.2019.1603378.

(31) Eadie, E.; Barnard, I. M. R.; Ibbotson, S. H.; Wood, K. Extreme Exposure to Filtered Far-UVC: A Case Study †. Photochem. Photobiol. 2021, 97 (3), 527–531. 10.1111/php.13385.

(32) ACGIH. Documentation on Threshold Limit Values and Biological Exposure Indices—Ultraviolet., 2022.

(33) Buonanno, M.; Welch, D.; Shuryak, I.; Brenner, D. J. Far-UVC Light (222 Nm) Efficiently and Safely Inactivates Airborne Human Coronaviruses. Sci. Rep. 2020, 10 (1), 10285. 10.1038/s41598-020-67211-2.

(34) Welch, D.; Buonanno, M.; Grilj, V.; Shuryak, I.; Crickmore, C.; Bigelow, A. W.; Randers-Pehrson, G.; Johnson, G. W.; Brenner, D. J. Far-UVC Light: A New Tool to Control the Spread of Airborne-Mediated Microbial Diseases. Sci. Rep. 2018, 8 (1), 2752. 10.1038/s41598-018-21058-w.

(35) ACGIH. Documentation on Threshold Limit Values and Biological Exposure Indices—Ultraviolet., 2021.

(36) Schmalwieser, A. W.; Wallisch, S.; Diffey, B. A Library of Action Spectra for Erythema and Pigmentation. Photochem. Photobiol. Sci. 2012, 11 (2), 251–268. 10.1039/c1pp05271c.

(37) UV-C Photocarcinogenesis Risks From Germicidal Lamps; Commission Internationale de l’Eclairage: CIE Central Bureau, Vienna, Austria, 2010.

(38) Cole, C. A.; Davies, R. E.; Forbes, P. D.; D’Aloisio, L. C. COMPARISON OF ACTION SPECTRA FOR ACUTE CUTANEOUS RESPONSES TO ULTRAVIOLET RADIATION: MAN and ALBINO HAIRLESS MOUSE*. Photochem. Photobiol. 1983, 37 (6), 623–631. 10.1111/j.1751-1097.1983.tb04531.x.

(39) Bissett, DonaldL.; Hannonand, DanielP.; Orr, ThomasV. AN ANIMAL MODEL OF SOLAR-AGED SKIN: HISTOLOGICAL, PHYSICAL, and VISIBLE CHANGES IN UV-IRRADIATED HAIRLESS MOUSE SKIN*. Photochem. Photobiol. 1987, 46 (3), 367–378. 10.1111/j.1751-1097.1987.tb04783.x.

(40) Benavides, F.; Oberyszyn, T. M.; VanBuskirk, A. M.; Reeve, V. E.; Kusewitt, D. F. The Hairless Mouse in Skin Research. J. Dermatol. Sci. 2009, 53 (1), 10–18. 10.1016/j.jdermsci.2008.08.012.

(41) Yamano, N.; Kunisada, M.; Nishiaki-Sawada, A.; Ohashi, H.; Igarashi, T.; Nishigori, C. Evaluation of Acute Reactions on Mouse Skin Irradiated with 222 and 235 Nm UV-C. Photochem. Photobiol. 2021, 97 (4), 770–777. 10.1111/php.13384.

(42) Chaney, E. K.; Sliney, D. H. RE-EVALUATION OF THE ULTRAVIOLET HAZARD ACTION SPECTRUM—THE IMPACT OF SPECTRAL BANDWIDTH. Health Phys. 2005, 89 (4), 322. 10.1097/01.HP.0000164650.96261.9d.

(43) Schneider, C. A.; Rasband, W. S.; Eliceiri, K. W. NIH Image to ImageJ: 25 Years of Image Analysis. Nat. Methods 2012, 9 (7), 671–675. 10.1038/nmeth.2089.

(44) Sandby-Møller, J.; Poulsen, T.; Wulf, H. C. Epidermal Thickness at Different Body Sites: Relationship to Age, Gender, Pigmentation, Blood Content, Skin Type and Smoking Habits. Acta Derm. Venereol. 2003, 83 (6), 410–413. 10.1080/00015550310015419.

(45) Egawa, M.; Hirao, T.; Takahashi, M. In Vivo Estimation of Stratum Corneum Thickness from Water Concentration Profiles Obtained with Raman Spectroscopy. Acta Derm. Venereol. 2007, 87 (1), 4–8. 10.2340/00015555-0183.

(46) Konger, R. L.; Derr-Yellin, E.; Hojati, D.; Lutz, C.; Sundberg, J. P. Comparison of the Acute Ultraviolet Photoresponse in Congenic Albino Hairless C57BL/6J Mice Relative to Outbred SKH1 Hairless Mice. Exp. Dermatol. 2016, 25 (9), 688–693. 10.1111/exd.13034.

