## Supporting Information. for "Extending the Acute Skin Response Spectrum to Include the Far-UVC"

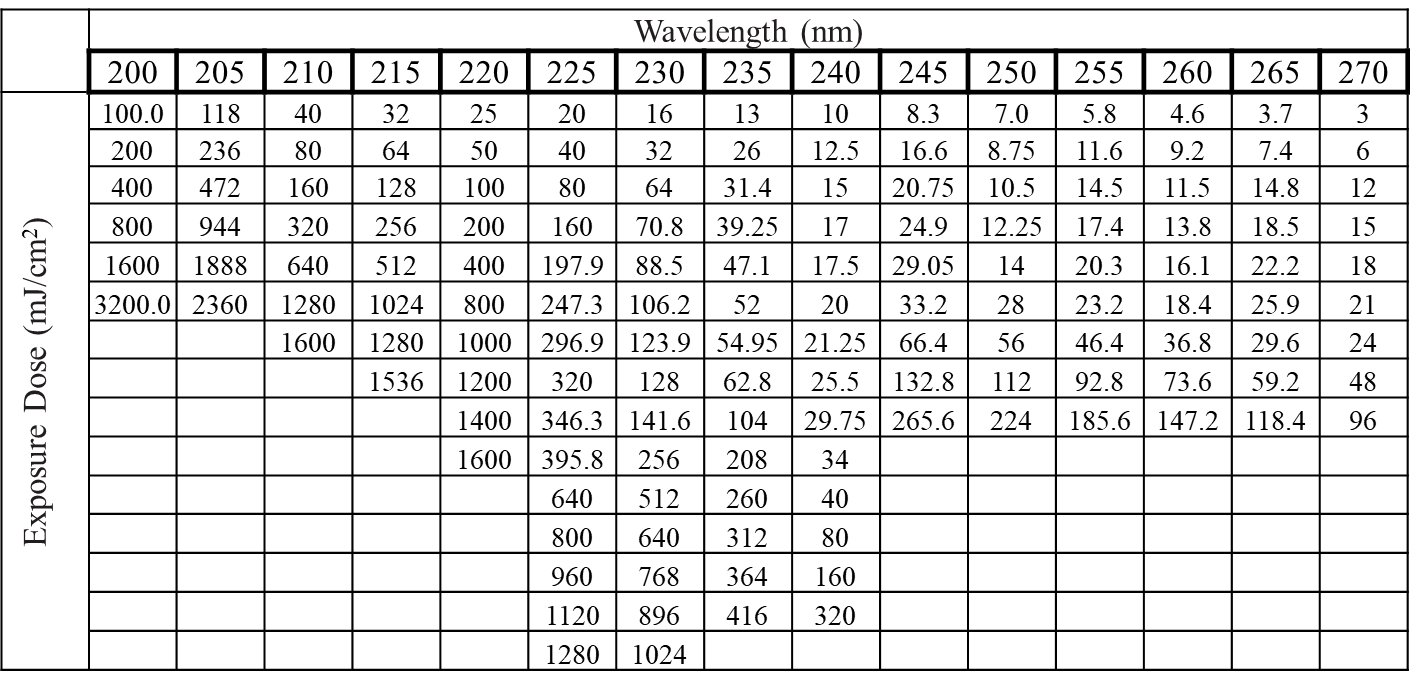

**Table S1.** Doses at each wavelength used to expose mice during the two stages of the study. The lowest exposure doses at each wavelength were based on the current 8-hour exposure limits as defined by ICNIRP,^1^ followed by increments of 2x, 4x, 8x, 16x, and 32x of these limits during the first stage of the study. The remaining of the exposure doses used at each wavelength were based on the estimated safe dose established during the initial stage of the process as well as 1.25x, 1.5x, 1.75x, and 2x this estimated safe dose.

**Stratum Corneum and Epidermis Thickness Measurements**

*Materials and Methods.* 72 h after exposure, mice were euthanized by CO2 asphyxiation and dorsal skin was harvested and immediately fixed in 10 % formalin (Epredia, Kalamazoo, MI) overnight at room temperature. Three randomly selected unirradiated skin tissues for each section of the grid were selected. Thickness measurements were performed by means of the Image-Pro Plus 6.0 software (Media Cybernetics, Silver Springs, MD) on hematoxylin-eosin stained 5 mm tissue sections. For the selected tissues, multiple randomly selected fields of view were analyzed across the tissues and the thickness of the stratum corneum and epidermis were collected following our previous mouse skin study.^1^ A one-way ANOVA test was used to statistically compare the thickness of the stratum corneum and epidermis of each dorsal section. Measurements represent the average ± standard deviation.

*Results.* The stratum corneum and epidermis on the dorsal side of hairless mice skin was found to have an average thickness of 4.70 ± 0.88 µm and 20.86 ± 5.52 µm, respectively (Table S2). Following a one-way ANOVA test, no statistical differences were observed across the various sections of the dorsal region that were irradiated with UVC light in this study.

| **Grid Section** | **Stratum Corneum Thickness (µm)** | **Epidermis**  **Thickness (µm)** |
| --- | --- | --- |
| 1 | 4.69 ± 1.15 | 20.19 ± 4.99 |
| 2 | 4.92 ± 1.00 | 20.33 ± 4.27 |
| 3 | 4.95 ± 0.67 | 22.54 ± 7.92 |
| 4 | 4.57 ± 0.68 | 21.60 ± 4.96 |
| 5 | 4.71 ± 0.89 | 20.69 ± 4.69 |
| 6 | 4.65 ± 0.93 | 21.95 ± 4.57 |
| 7 | 4.53 ± 0.88 | 20.34 ± 5.96 |
| 8 | 4.63 ± 0.65 | 19.26 ± 5.22 |

**Table S2.** Average thickness of the (a) stratum corneum and (b) epidermis were determined based on histological analysis of three skin tissues per section of the grid drawn on the dorsal side of the mice. Overall, the average thickness of the stratum corneum and epidermis throughout the dorsal side of SKH-1 albino hairless mice was found to be 4.70 ± 0.88 µm and 20.86 ± 5.52 µm, respectively.

**References.**

1.	The International Commission on Non-Ionizing Radiation Protection. GUIDELINES ON LIMITS OF EXPOSURE TO ULTRAVIOLET RADIATION OF WAVELENGTHS BETWEEN 180 nm AND 400 nm (INCOHERENT OPTICAL RADIATION). *Health Physics* **87**, 171–86 (2004).

2.	Buonanno, M. *et al.* 207-nm UV Light—A Promising Tool for Safe Low-Cost Reduction of Surgical Site Infections. II: In-Vivo Safety Studies. *PLoS One* **11**, e0138418 (2016).
